# Towards Robust Automated Mouse Scratching Behavior Detection for Itch Research

**DOI:** 10.64898/2026.10.12.758682

**Authors:** Shu Yang, Anyu Pan, Hyowon Shin, Hongzhuo Chen, Zhao Zhang, Azim Shad, William Huang, Yang Liu, Wesley Liu, Helen Luo, Chentianye Xu, Jiong Chen, Min Xu, Wenqin Luo, Huasheng Yu, Li Shen

**Affiliations:** University of Pennsylvania, Philadelphia, USA; Carnegie Mellon University, Pittsburgh, USA

**Keywords:** mouse behavior, itch research, video foundation models, multimodal large language models, transfer learning

## Abstract

Quantification of mouse scratching behavior is essential for preclinical itch research. However, manual video annotation is labor-intensive, and automated detection remains challenging in terms of accuracy and generalizability. We present Scratch-AID+, a computational framework dedicated to automated mouse scratching detection, including a diverse and progressive set of model families and learning strategies: classic ResNet-based detectors, task-adaptive fine-tuning of video foundation models, and in-context learning of advanced multimodal large language models. To simulate real-world use cases, we benchmark on a public dataset containing annotated video recordings from a source laboratory with additional cross-laboratory videos for independent testing, and we explore different data preprocessing and training configurations. We find that classic models suffer severe performance degradation on external data under our realistic setting with sparse scratching events. In contrast, fine-tuned video foundation models substantially improve the generalizability, achieving significantly better detection performance. Furthermore, frozen multimodal large language models demonstrate promising out-of-the-box ability using only few-shot demonstrations. We also report the effects of different video preprocessing strategies and ablations. Our findings highlight the improved performance and generalizability of the adapted foundation models for automated scratching behavior analysis and shed light on the real-world deployment of such detectors. Our code is publicly available at github.com/PennShenLab/scratch-aid-plus.

## I. Introduction

Reliable measurement of scratching behavior is critical for studying the biological mechanisms of itch and for assessing the efficacy of potential antipruritic treatments [1]. Mouse models are used extensively in this field, and the frequency of scratching is commonly considered as the principal behavioral measure of itch severity [2], [3]. Traditionally, mouse scratching is quantified by manually reviewing recorded videos, a process that is time-consuming, difficult to scale, and vulnerable to differences between observers. These limitations make manual analysis especially challenging for large genetic, pharmacological, or therapeutic screening studies. Several automated approaches have therefore been developed, including systems based on acoustic signals, electromagnetic sensing, and machine-learning or deep-learning analysis of video recordings [4]–[7]. Despite this progress, automated rodent-scratching detection methods have not been widely adopted across the itch research field.

A major barrier to broader adoption is model generalization: a model trained in one laboratory or recording setting may not perform well when applied to videos from another laboratory. For example, one of the best available systems for mouse scratching detection, Scratch-AID [8], uses a convolutional recurrent neural network (CRNN) framework to achieve high accuracy within source environments. However, its cross-laboratory robustness requires further investigation, since acquisition conditions and the distribution of scratching events can differ. Recently, large-scale vision-language models and benchmarks have begun to emerge, providing specialized resources for rodent behavior analysis [9]–[11]. Despite these advances, the cross-laboratory generalization of foundation models and classic detectors for the specific sparse signal of mouse scratching has not been systematically studied.

In the general computer vision domain, pretrained video foundation models (vFMs), such as VideoMAE [12], InternVideo-Next [13], and VideoSwin Transformer [14], often provide complementary representations of appearance and motion. Large-scale pretraining supplies reusable spatiotemporal features that can be adapted through task-specific fine-tuning. Meanwhile, multimodal large language models (MLLMs), such as Qwen [15] and Gemini [16], offer a different adaptation mechanism: learning from visual input and labeled in-context examples without updating model weights.

To address this generalization gap, we present Scratch-AID+, extending Scratch-AID to provide a much improved framework dedicated to mouse scratching detection that utilizes and compares ResNet-based baselines, adapted vFMs, and frozen MLLMs. These distinct approaches reuse pretrained knowledge in different ways: the baseline models combine frame-level features derived from ResNet with task-trained temporal components, vFMs transfer spatiotemporal representations through full or parameter-efficient fine-tuning, and MLLMs condition predictions on labeled in-context examples without task-specific weight updates. In addition, we examine diverse cropping and transformation approaches to systematically assess how video preprocessing affects cross-laboratory performance for mouse scratching detection. We use the publicly available Scratch-AID dataset for model training, validation, and in-lab testing, and we also obtain additional data from four external laboratories (videos recorded with the same device as Scratch-AID) for held-out evaluation. This study can help to lay the foundation for robust, generalizable, and automated mouse scratch detection for itch research.

## II. Methods

### A. Data and Processing

Following Scratch-AID [8], Scratch-AID+ formulates the mouse scratching detection task as binary video-clip classification (Fig. 1): class 1 denotes scratching and class 0 denotes non-scratching. The source in-lab dataset comprises 40 20-minute videos (frame rate=30 fps) from 10 C57BL/6J mice (5 male and 5 female) following pharmacological induction of scratching. 32 videos from 8 mice are used for training, and 8 videos from 2 mice (different from the 8 mice in the training videos) are used for evaluation. The held-out external dataset comprises 16 additional 20-minute videos recorded from 16 mice across 4 different labs. These videos were manually annotated frame-by-frame using the same standard as described for Scratch-AID [8].

**Fig. 1:**
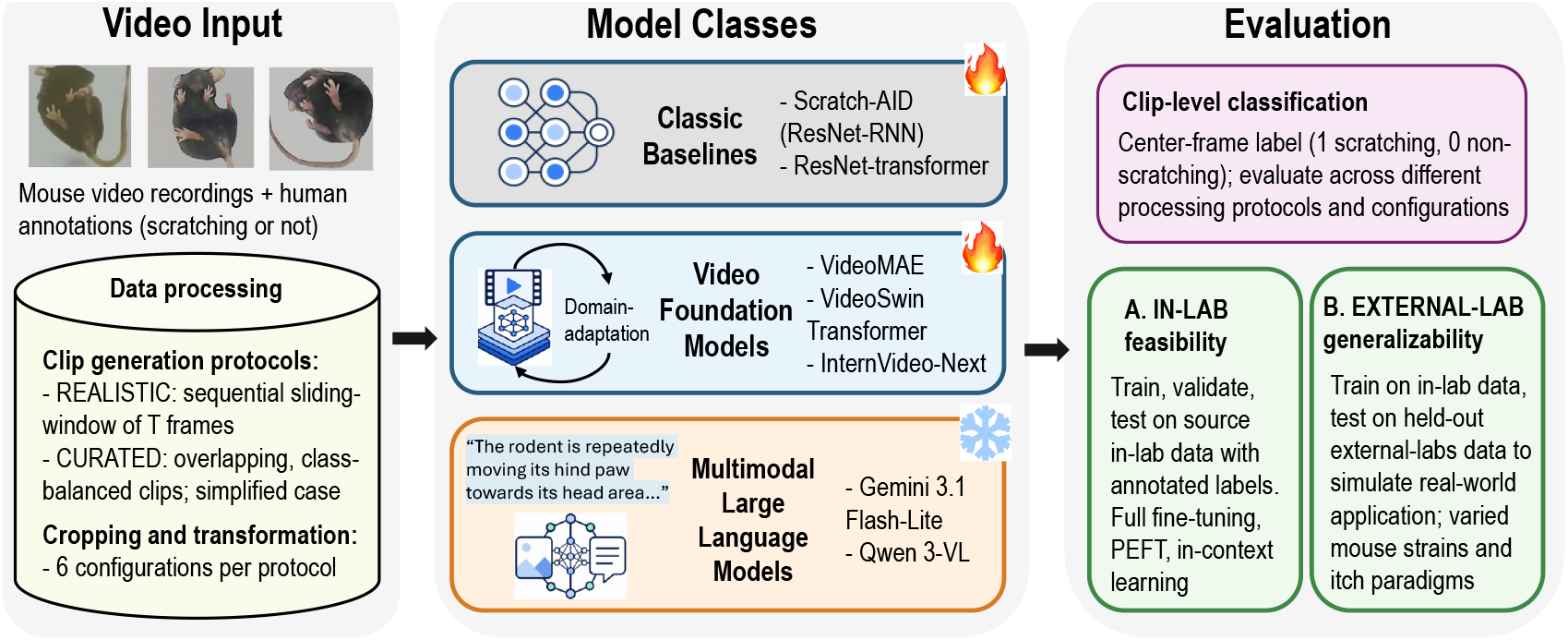
Overview of the Scratch-AID+ workflow. Classic baselines and pretrained video foundation models are adapted using source in-lab data for further supervised training, whereas multimodal LLMs remain frozen and utilize in-context learning. Scratching-class F1 is reported separately for in-lab and external tests across the REALISTIC and CURATED protocols.

From the original long video recordings, we extract short clip segments for models to process. Let *x* = (*x*_1_, …, *x*_*T*_) be a clip of *T* frames with binary annotations *a*_*t*_ ∈ {0, 1}. For odd *T*, the clip label *y* is determined by the center frame annotation *y* = *a*_(*T* +1)*/*2_. We then employ two distinct evaluation protocols: **1) REALISTIC**, our *main* evaluation protocol, uses clips extracted sequentially via non-overlapping sliding windows of length *T* to simulate continuous screening where scratching events are naturally sparse. **2) CURATED** is a class-balanced set of overlapping clips, representing a simplified and idealized task setting. Positive clips contain at least (*T* + 1)*/*2 consecutive scratching frames, while negatives are sampled at least 45 frames away from any scratching event. We use *T* = 29 frames (approximately 1 second), balancing temporal context and GPU memory constraints. In general, this clip-based formulation enables annotation of any frame in the original recording, with the target frame centered in the clip and flanking frames providing temporal context.

To facilitate robustness against mouse/recording variability, we apply six video preprocessing configurations to each protocol, generated by two cropping strategies and three color transformations. Spatial crops include *Per-Frame* mode, which independently selects the lowest-mean-intensity 256 ×256 window in each frame, and *Motion-Gradient* mode, which selects one window for the entire clip using normalized motion and gradient scores weighted 0.6 and 0.4. Color transformations include *None* (original RGB), *Gray* (grayscale), and *Color Jitter* (brightness, contrast, saturation, and hue perturbations with parameters (0.4, 0.4, 0.4, 0.1)). We denote the resulting clip by 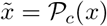 for configuration *c* ∈ {1, …, 6}.

### B. Models

Given a processed clip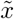, each model predicts a binary label *ŷ* ∈ {0, 1}. Here, Scratch-AID+ includes three model classes (Fig. 1), representing both conventional baselines and recent paradigms in video foundation models and generative AI, for automated scratching detection task:

#### Classic baselines

We benchmark the original Scratch-AID [8], which utilizes a convolutional recurrent neural network (CRNN) architecture based on ResNet-18, and we also extend it to a ResNet-transformer baseline, replacing the RNN with a temporal transformer:

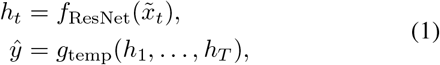

where *g*_temp_ includes temporal modeling by RNN or transformer, combining pretrained ResNet-18 frame-level features with a task-trained classifier.

#### Video foundation models (vFMs)

To assess transfer from general-purpose video representations, we adapt vFMs pretrained on large-scale video datasets (e.g., Kinetics-400 [17]). Three top-ranked models, VideoMAE [12], VideoSwin transformer [14], and InternVideo-Next [13], are included here for domain adaptation with full or parameter-efficient fine-tuning (PEFT). A video encoder *f*_vid_ and a pooling/classification head *g*_cls_ jointly process spatial and temporal information:

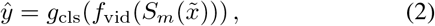

where *S*_*m*_ denotes specific temporal sampling and resizing for model *m*.

#### Multimodal large language models (MLLMs)

Gemini3.1 Flash-Lite [16] and Qwen3-VL-32B-Instruct [15] are adopted, representing the class of advanced LLM-based methods. With frozen model weights *θ*_0_, task prompt *I*, and *k* labeled training clips 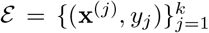, they rely on few-shot in-context learning (ICL):

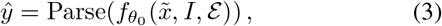

where Parse extracts the generated class label.

### C. Training and Evaluation

Here, we present the general training and evaluation workflows. Detailed model implementations, input pipelines, and ICL prompts are available in our Github repo.

#### Training settings

All models use the fixed source training/validation/test partitions of Scratch-AID [8] (Table I). To prevent data leakage, all clips from a single recording remain in the same data split. External clips serve strictly as independent held-out sets to simulate real-world cross-laboratory deployment without any target-domain adaptation. All trainable models (classic baselines and vFMs) are optimized via binary cross-entropy loss. Hyperparameters such as learning rate, weight decay, dropout, LoRA rank, and alpha are tuned on validation sets using Optuna [18]; full search ranges and inference settings are available in the public Github repo. We explored full fine-tuning vs. PEFT for the trainable models and different numbers of shots (0,2,4) in ICL for frozen MLLMs in preliminary analyses to save computation costs and report the selected setting for each model. Scratch-AID, ResNet-transformer, VideoMAE, InternVideo-next, and VideoSwin are fine-tuned for the scratching task (e.g., VideoMAE starts from Kinetics-400-pretrained weights and adopts full fine-tuning, whereas InternVideo-Next uses LoRA [19] due to GPU memory constraints). For MLLM in-context learning, class-balanced *k*-shot demonstrations are retrieved from the training set via cosine similarity-based nearest-neighbor search over training video embeddings. To limit inference costs, MLLMs experiments are conducted on three independent random subsamplings per protocol/configuration, each with 200 training and 200 test clips (100 per class).

**TABLE I:** Dataset partitions and clip counts. Parentheses give scratching-positive counts.

| Partition | REALISTIC | CURATED |
|---|---|---|
| Training | 39,712 (9,020) | 95,582 (47,791) |
| Validation | 4,964 (908) | 9,344 (4,672) |
| In-lab test | 4,964 (980) | 9,972 (4,986) |
| External test | 18,616 (1,428) | 14,228 (7,114) |

Note: The train/validation/test partitions follow scratch-AID [8]. Each row applies to all six video preprocessing configurations.

#### Implementation environment

The Gemini model is accessed via Google’s official API, while all the other models run locally on a high-performance cluster using NVIDIA A100 (80 GB vRAM) GPUs with 160 GB main memory per node. Table II reports approximate model sizes (estimate of model complexity). Due to the large number of experimental configurations, we executed several hundred cluster jobs in total with a combined over *5,000* GPU hours.

**TABLE II:** Scratching-class F1 (%) averaged over six preprocessing configurations.

|  |  | REALISTIC |  | CURATED |  |
|---|---|---|---|---|---|
| Model | # Params. | In-lab | External | In-lab | External |
| *A. Trainable models* |  |  |  |  |  |
| Scratch-AID | 12M | 95.98 | 66.31 | 99.67 | 85.40 |
| ResNet-transformer | 12M | 96.60 | 71.20 | 99.68 | 85.87 |
| VideoMAE | 304M | 97.00 | **82.49** | 99.92 | 95.04 |
| VideoSwin | 88M | 97.03 | 78.12 | 99.76 | 93.00 |
| InternVideo-Next | 94M | 96.99 | 81.19 | 99.86 | **95.26** |
| *B. Frozen MLLMs with four-shot ICL* |  |  |  |  |  |
| Gemini3.1 Flash-Lite | Undisclosed | 77.50 | 75.07 | 77.07 | 73.19 |
| Qwen3-VL | 32B | 80.15 | 72.43 | 78.41 | 76.26 |

Parameter counts are approximate. Trainable models use full test sets. MLLMs use three random, class-balanced subsamples per configuration, as described in II-C; MLLM scores reported here are averaged first across random repeats and then configurations. Bold marks the highest external F1 among trainable models within each protocol.

#### Evaluation metrics

For evaluation, scratching-class (class-1) F1 score is the primary metric; accuracy is secondary and reported only in the supplementary Table S1 on our Github repo. Similarly, for each model, protocol, and test domain (in-lab, external), Section III reports the mean F1 across the configurations 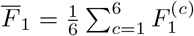, with a detailed breakdown in Table S1.

## III. Results

### A. Within- and Cross-Laboratory Performance

Our large-scale experiments enable systematic evaluation across model classes and video processing strategies while maintaining strict separation of training, test, and validation data. Table II reports mean scratching-class F1 across six preprocessing configurations (full results in Table S1). We focus on the REALISTIC protocol, with CURATED serving as a complementary benchmark.

On in-lab REALISTIC clips, trainable models achieved similar F1 scores (95.98 - 97.03%). Differences were larger on external clips (Table II and Fig. 2A): VideoMAE reached 82.49%, followed by InternVideo-Next (81.19%) and VideoSwin (78.12%), compared with Scratch-AID (66.31%) and ResNet-transformer (71.20%). In particular, VideoMAE exceeded Scratch-AID by 16.18 percentage points. Indeed, all three vFMs achieved higher mean external REALISTIC F1 than both ResNet-based classic baselines, and they exceeded Scratch-AID in each configuration. On CURATED clips, in-lab F1 ranged from 99.67% to 99.92%; for external F1, InternVideo-Next (95.26%) slightly exceeded VideoMAE (95.04%), reversing their REALISTIC ordering.

**Fig. 2:**
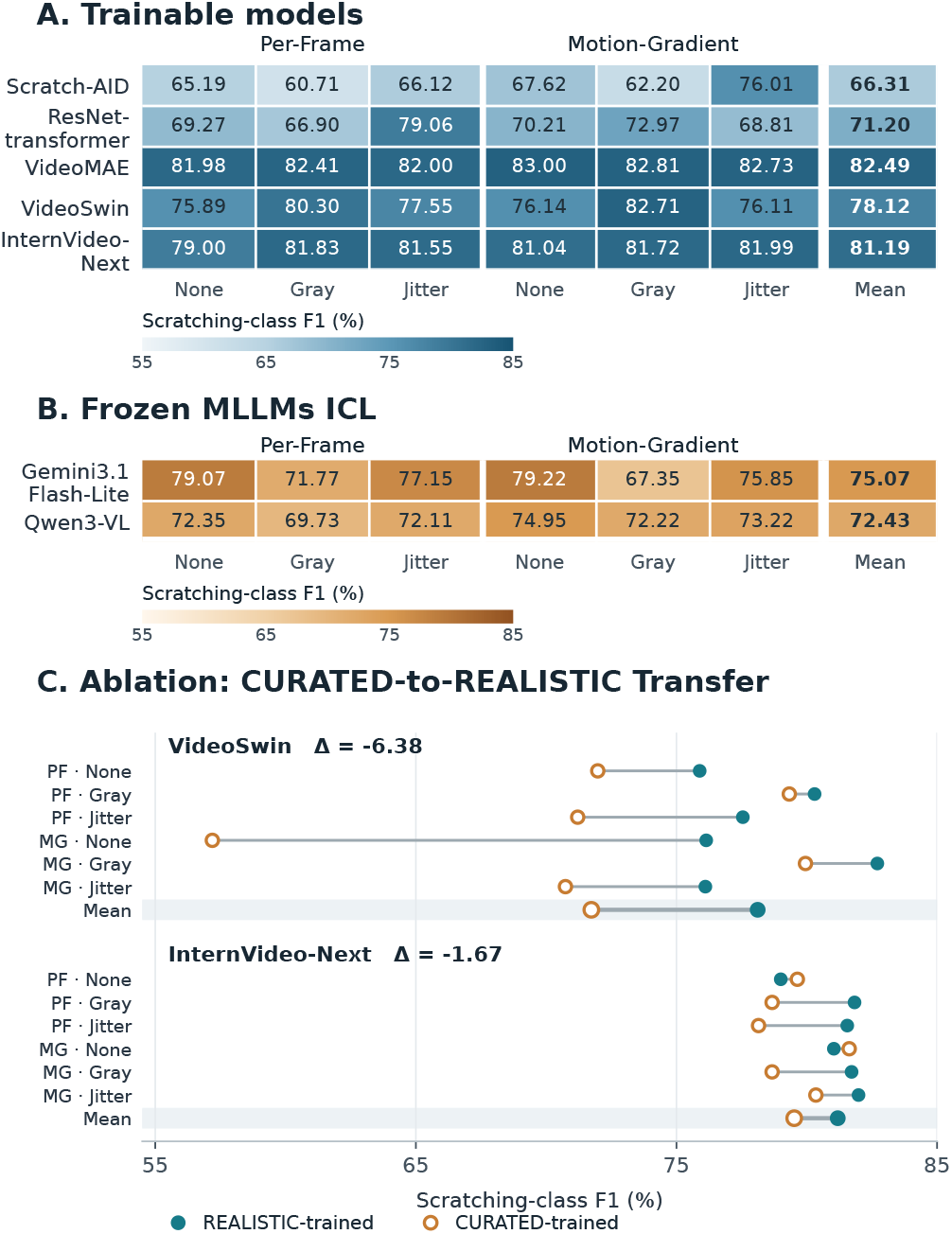
External realistic evaluation. (A) Per-configuration F1 heatmap for trainable models. (B) F1 heatmap for frozen MLLMs. (C) Ablation study of CURATED-trained check-points evaluated on the REALISTIC test set. PF/MG: Per-Frame/Motion-Gradient cropping; Jitter: color jitter.

#### Precision & recall

It is worth noting that only 7.67% of external REALISTIC clips were positive due to the sparse nature of the scratching events, making accuracy metric insufficient since an always-negative classifier would achieve 92.33% accuracy. Thus, we put the accuracy results in supplementary Table S1 for completeness and examined the confusion matrix for more details. Under the *Per-Frame None* configuration, VideoSwin yielded True Positive/False Positive/True Negative/False Negative = 1,086/348/16,840/342, with precision=75.73% & recall=76.05% on the 18,616 external test clips; similarly, InternVideo-Next yielded 1,125/295/16,893/303, with precision=79.23% & recall=78.78%. Across six configurations, mean precision & recall are 74.51 & 82.55% for VideoSwin and 79.41 & 83.09% for InternVideo-Next, respectively.

#### Matched-subset comparison

On balanced external REALISTIC-derived subsets as described in Section II-C, frozen Gemini and Qwen achieved mean F1 of 75.07% and 72.43% across six configurations (Fig. 2B) and three random repeats, respectively (Table II). To make a fair comparison, VideoSwin and InternVideo-Next were evaluated on the same subsets, yielding 89.49% and 90.48% mean F1. Despite lacking task-specific training to further improve the performance, these cutting-edge MLLMs exhibited promising ICL ability and small cross-laboratory degradation.

### B. Ablation Analysis

#### Protocol transfer

As an ablation, we reused the same CURATED-trained model checkpoints of VideoSwin and InternVideo-Next on REALISTIC clips without retraining, comparing them with checkpoints trained and selected under REALISTIC (Fig. 2C). VideoSwin and InternVideo-Next achieved F1 of 71.73% and 79.52%, respectively, below the 78.12% and 81.19% obtained by checkpoints trained directly under REALISTIC. Thus, the benefits of class-balanced training and clean frame composition of CURATED clips do not seem to carry over to the real-world sparse-scratching-event evaluation in REALISTIC.

#### Preprocessing sensitivity

External REALISTIC F1 varied little for VideoMAE (81.98-83.00%) and InternVideo-Next (79.00-81.99%), whereas Scratch-AID spanned 15.30 percentage points (Fig. 2A). Under *Per-Frame* cropping, *grayscale* increased VideoSwin F1 from 75.89% to 80.30% but reduced Scratch-AID from 65.19% to 60.71%; both effects had the same direction under *Motion-Gradient* cropping. Preprocessing benefits seem to be highly model dependent.

## IV. Discussion and Conclusion

Scratch-AID+ extends Scratch-AID through diverse strategies for improving mouse scratching detection, inspired by the recent advancements and paradigm shift in machine learning. It includes transformer temporal modeling, adapted video foundation models, and advanced multimodal LLMs. The systematic benchmarking establishes an essential foundation for the robust, cross-laboratory deployment of animal behavior analysis in the era of generative AI.

Our results highlight the potential of adapted vFMs to improve automated scratching detection across laboratories. Although trainable models achieved similar performance on in-lab recordings, vFMs consistently achieved higher external F1 scores than the ResNet-based baselines. This pattern is consistent with the possibility that large-scale video pretraining provides transferable representations of appearance and motion that mitigate overfitting and support generalization beyond the source environment. Moreover, the external evaluation included four laboratories using the same recording device as the source dataset; broader robustness across camera systems and acquisition conditions remains to be established. Despite their improved generalization in our evaluation, vFMs still exhibited a performance gap between within-laboratory and external-laboratory recordings. Addressing this gap will likely require advances in both model development and video acquisition.

Pretraining on larger and more diverse animal behavior datasets spanning multiple laboratories, recording conditions, and behavioral contexts may improve the transferability of learned representations. In parallel, systematically identifying sources of distribution shift, such as differences in lighting, resolution, and frame rate, could guide acquisition standardization and targeted mitigation strategies. Together, these approaches may improve the reliability of automated scratching detection across laboratories.

Frozen MLLMs also demonstrated promising scratching recognition using only few-shot in-context demonstrations and no task-specific weight updates. On matched external subsets, their performance remained below that of the fine-tuned vFMs, but their ability to recognize this specialized behavior suggests potential for reducing the amount of task-specific annotation and model training required. Further improvements in accuracy, through optimized selection and presentation of labeled demonstrations and advances in the underlying MLLMs, will be critical for practical use and deployment.

Preprocessing affected detection performance in a model-dependent manner. For example, grayscale conversion improved VideoSwin performance but reduced Scratch-AID performance, whereas VideoMAE and InternVideo-Next were relatively stable across the evaluated configurations. These differences suggest that preprocessing may alter the balance between appearance and motion cues used by each model. Cropping can also influence animal visibility and temporal consistency. These mechanisms remain hypotheses, but the observed variation indicates that preprocessing should be validated jointly with the selected model rather than assumed to provide a universal benefit.

Several limitations should be considered for future work. (1) In our study, recorded mean processing times were 0.23 s/clip for InternVideo-Next’s test pipeline, 5.08 s/clip for Gemini API calls, and 8.98 s/clip for Qwen inference. Differences in model access, test views, and timing boundaries preclude a controlled comparison of speed or monetary cost. Nevertheless, real-world deployment decisions should consider available computing resources, training and inference demands, and reliance on local hardware or external APIs alongside recognition accuracy. (2) External validation included 16 mice from four laboratories using the same recording device as the source dataset; therefore, our findings support cross-laboratory generalization under the evaluated acquisition conditions, while robustness to broader hardware and recording differences remains untested. Laboratory-specific evaluations with more videos based on independent animals or recordings are needed, because average performance across different laboratories does not reflect performance reproducibility and variability in different conditions. (3) Although current matched-subset analyses enabled comparisons between model families, MLLM evaluation primarily used small, randomly sampled, and approximately balanced subsets because of cost constraints and does not establish performance under naturally sparse scratching prevalence. (4) Finally, center-frame classification of short clips can be challenging near behavioral transitions, and clip-level F1 does not directly establish accurate video-wise scratching-event counts or duration estimates. Further validation of temporal localization, false positives per recording hour, and deployment costs will be important for routine experimental use.

## Acknowledgment

We thank the laboratories of Drs. Xingzhong Dong, David Ginty, and Xintong Dong for their support and assistance with video recording for external validation. This work was supported in part by seed funding from the PSOM AI2D Center at Penn (L.S.) and NIH grant R01-NS-131209 (W.L.).

